# Purification and Immunogenicity of Nipah Virus-Like Particles from Insect Cells

**DOI:** 10.1101/2025.08.21.671198

**Authors:** Urban Bezeljak, Alexander Jerman, Tina Kobal, Martina Lokar Kosmač, Elfi Birsa, Marko Kolenc, Dániel Déri, Bernadett Pályi, Zoltán Kis, Matjaž Peterka

**Affiliations:** Sferogen d.o.o., Ajdovščina, Slovenia; COBIK, Ajdovščina, Slovenia; Institute of Microbiology and Immunology, Faculty of Medicine, University of Ljubljana, Ljubljana, Slovenia; National Biosafety Laboratory, National Center for Public Health and Pharmacy, Budapest, Hungary; European Research Infrastructure on Highly Pathogenic Agents (ERINHA-AISBL), Brussels, Belgium

**Keywords:** virus-like particles, Nipah virus, baculovirus expression, monolith chromatography, insect cell culture, vaccine development, pandemic preparedness

## Abstract

Nipah virus (NiV) is an emerging high-fatality zoonotic threat lacking approved vaccines. Current virus-like particle (VLP) production methods rely on costly mammalian cell systems and non-scalable ultracentrifugation purification. We developed an efficient production system for enveloped NiVLPs by co-expressing structural proteins F, G, and M using a baculovirus expression vector system in Sf9 cells. A novel multi-step chromatographic purification process was established using monolith convective media, integrating steric exclusion chromatography with sequential cation and anion exchange steps. Purified NiVLPs were characterized by nanoparticle tracking analysis and transmission electron microscopy, then evaluated for immunogenicity in Syrian golden hamsters. The optimized process yielded enveloped particles of approximately 100-120 nm that morphologically resemble native NiV virions. A single 25 μg NiVLP dose induced robust systemic anti-NiV G IgG responses within 14 days, demonstrating rapid immunogenicity suitable for outbreak response. However, neutralizing antibody titers against NiV remained limited compared to total IgG responses. This study establishes the first chromatography-based manufacturing platform for morphologically correct NiVLPs from insect cells. A deeper understanding of the immunity generated is needed to support their potential as a rapidly deployable vaccine platform against NiV.

## Background

Nipah virus (NiV) is a zoonotic paramyxovirus, which was first identified in 1998 during an outbreak in Malaysia and Singapore. Since its emergence, NiV has reappeared almost annually in outbreaks in Bangladesh and India, where it continues to cause severe disease (Khan *et al*., 2024). NiV is a risk group 4 pathogen and is associated with severe respiratory disease and encephalitis, resulting in fatality rates ranging from 40-100% (Amaya and Broder, 2020). The natural reservoir of NiV is fruit bats of the *Pteropodidae* family. Transmission to humans can occur directly, or via intermediate hosts such as pigs and horses (Eaton *et al*., 2006). Human-to-human transmission has also been documented, notably by the respiratory route.

The lack of an approved Nipah vaccine makes NiV a high-priority pathogen with pandemic potential (Moore *et al*., 2024). The development of diverse vaccine platforms offering broad and lasting protection is therefore urgently needed. Various immunization strategies have been explored, including mRNA (Lo *et al*., 2020; Loomis *et al*., 2021; Pedrera *et al*., 2024), DNA (Lu *et al*., 2023), viral vector (Yoneda *et al*., 2013; de Wit *et al*., 2022; Foster *et al*., 2022; Ithinji *et al*., 2022; van Doremalen *et al*., 2022), protein subunit (Mungall *et al*., 2006; Pallister *et al*., 2013; Loomis *et al*., 2020; Geisbert *et al*., 2021; Gao *et al*., 2022), and virus-like particles (VLPs) (Walpita *et al*., 2011, 2017; Welch *et al*., 2023).

The surface NiV receptor-binding glycoprotein (G) is the major immunogen that generates neutralizing antibodies and is included in all vaccine candidates that progressed to clinical trials (Kim *et al*., 2025). NiV G binds to conserved ephrin-B2/B3 receptors (Xu *et al*., 2008; Wong *et al*., 2021; Larsen *et al*., 2025), which leads to membrane fusion mediated by the NiV fusion glycoprotein (F). Likewise, NiV F can also generate a neutralizing immune response (Avanzato *et al*., 2019; Dang *et al*., 2021), albeit lower than NiV G (Wit *et al*., 2023). The matrix (M) and nucleocapsid (N) are the remaining structural proteins that are involved with virion budding and RNA genome encapsidation, respectively (Watkinson and Lee, 2016; Ker *et al*., 2021).

VLPs are a proven and safe subunit vaccine platform (Noad and Roy, 2003; Donaldson *et al*., 2018; Nooraei *et al*., 2021) that are non-infectious as they do not contain any genetic material. VLPs are self-assembled nanoparticles composed of viral structural proteins that closely resemble native pathogens and have the potential to present all key NiV immunogens. Nipah VLPs (NiVLPs) composed of NiV M, F, and G proteins have been previously explored using mammalian cell expression systems (Walpita *et al*., 2011). Co-expression of these proteins was shown to orchestrate the formation and budding of NiV VLPs that appeared structurally similar to authentic NiV virions and provide protection against viral challenge (Walpita *et al*., 2017). However, so far NiVLP purification strategy relied on ultracentrifugation. This approach is not suitable for vaccine production scale-up (Effio and Hubbuch, 2015) and does not resolve the enveloped VLPs from host cell contaminants (Margine *et al*., 2012; Minh and Kamen, 2021). Instead, a chromatography-based downstream process for NiVLPs needs to be established to support advanced nanoparticle vaccine development (Morenweiser, 2005; Gagnon, 2009; Nestola *et al*., 2015).

In this work, we aimed to address purification challenges and develop an alternative production system. We describe the generation of enveloped NiVLPs incorporating key structural proteins (G, F, M) in an insect cell expression system. We developed and optimized a scalable, multi-step purification strategy employing convective interaction media (monoliths). This process integrates steric exclusion chromatography (SXC) with subsequent cation (CEX) and anion (AEX) exchange steps on monolith columns. The purified NiVLPs were characterized by nanoparticle tracking analysis (NTA) and transmission electron microscopy (TEM). Furthermore, these particles elicited humoral immune responses in Syrian golden hamster model. This outlined downstream process, leveraging the benefits of convective media, provides a robust and scalable platform adaptable for purifying similar enveloped nanoparticles for diverse applications, including VLP vaccines against emerging viruses with pandemic potential.

## Results

NiVLPs can self-assemble from co-expressed structural proteins in mammalian cells (Walpita *et al*., 2011). However, this system presents challenges for cost-effective, large-scale vaccine production. Insect cells offer an alternative scalable expression system suitable for VLP production, as demonstrated for influenza (Smith *et al*., 2013) and coronaviruses (Mortola and Roy, 2004; Bezeljak *et al*., 2025). We therefore hypothesized that a baculovirus system could drive NiVLP assembly and secretion in insect cells. To test this, we constructed a single recombinant baculovirus (rBV) to co-express NiV M, F, and G proteins in Sf9 cells (Fig. 1a). Following 4 days of expression, culture supernatant was harvested via centrifugation. To capture nanoparticles, including potential NiVLPs, we used SXC principles using polyethylene glycol (PEG) for initial capture of nanoparticles from clarified supernatant (Lee *et al*., 2012). Unfiltered supernatant containing 6 % PEG6000 was applied to an 8 mL CIM OH column, facilitating binding of larger particles while removing smaller contaminants in flow-through. Bound particles were subsequently eluted with a decreasing PEG6000 gradient (Fig. 1b). Analysis of NiV G protein content served as an indicator of NiVLP presence in the collected fractions (Fig. 1c). The detection of NiV G across elution fractions suggests successful NiVLP assembly and secretion in insect cells. Notably, high PEG concentrations in E1 altered the migration of the NiV G on PAGE and western blot, resulting in a lower apparent molecular weight.

**Fig. 1.**
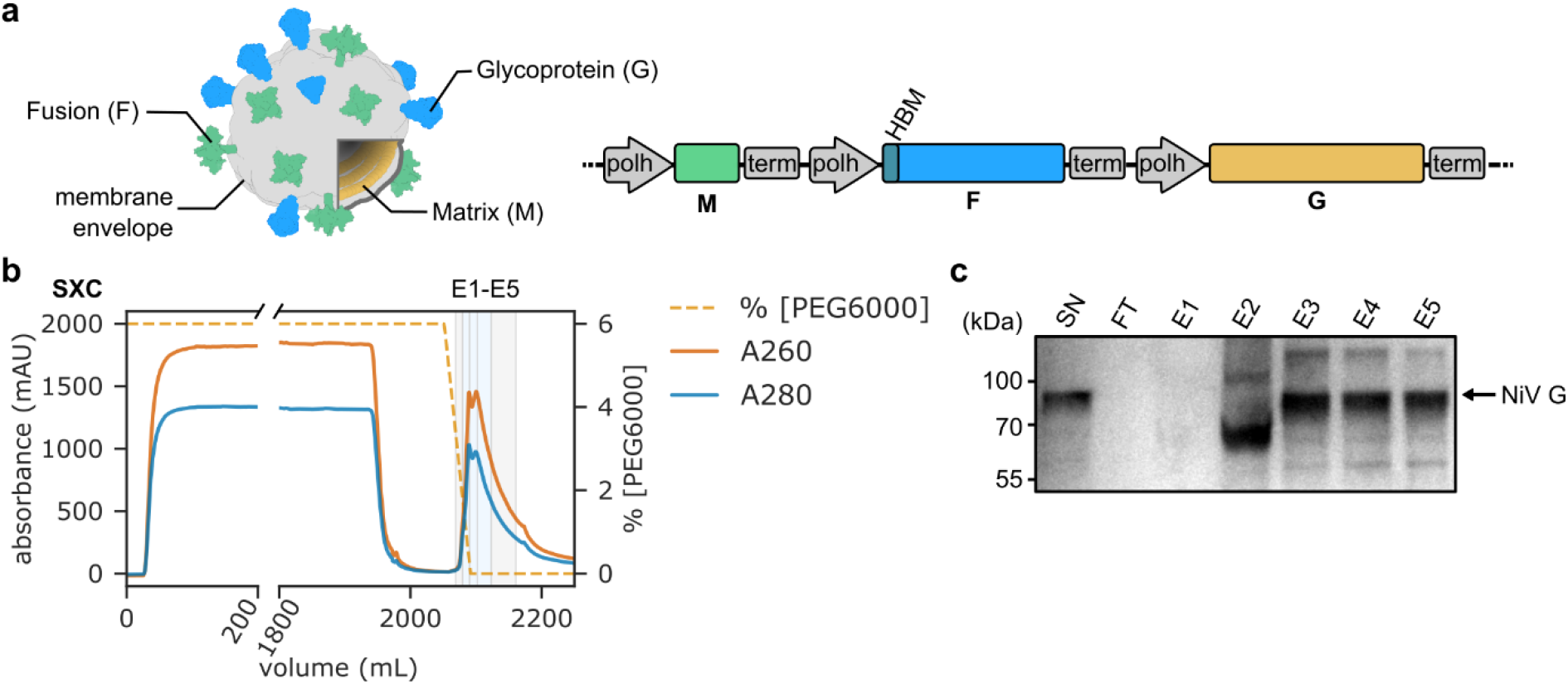
NiVLPs are captured with SXC. **(a)** Baculovirus expression cassette for the co-expression of NiV M, F and G structural proteins in Sf9 insect cells. HBM: honeybee melittin signal sequence; polh: polyhedrin promoter; term: SV40-terminator. **(b)** NiVLP capture step on CIM OH column. Expressed NiVLPs are bound to 8 mL CIM OH column in presence of 6 % PEG6000 from insect cell culture supernatant. Particles were released in descending [PEG6000] gradient. Shaded boxes denote elution fractions E1-5. Fractions E2-4, which were selected for further purification, are light blue. **(c)** Western blot analysis of capture chromatography step. NiV G protein is detected in E2-E5 elution fractions. SN: cell culture supernatant; FT: chromatography flow-through; E1-5: elution fractions.

We proceeded with purification of NiV G-containing fractions from the capture step using CEX chromatography. Diluted SXC elution fractions were loaded onto an 8 mL CIM SO3 column and eluted via a step gradient of ascending NaCl concentration (Fig. 2a). While NiV G was present in all fractions, the majority eluted in fraction E2 (15 to 33 mS/cm conductivity) (Fig. 2b). Finally, NiVLPs were further purified by AEX chromatography. The collected CIM SO3 E2 fraction was applied to a 4 mL CIM QA monolith, and elution was performed using a linear ascending salt gradient (Fig. 2c). Western blot analysis revealed NiV G eluted in a peak centered around the E2 fraction (Fig. 2d). Finally, buffer was exchanged into PBS with 15 % sucrose and 0.05 % polysorbate-80 using gel filtration group separation.

**Fig. 2.**
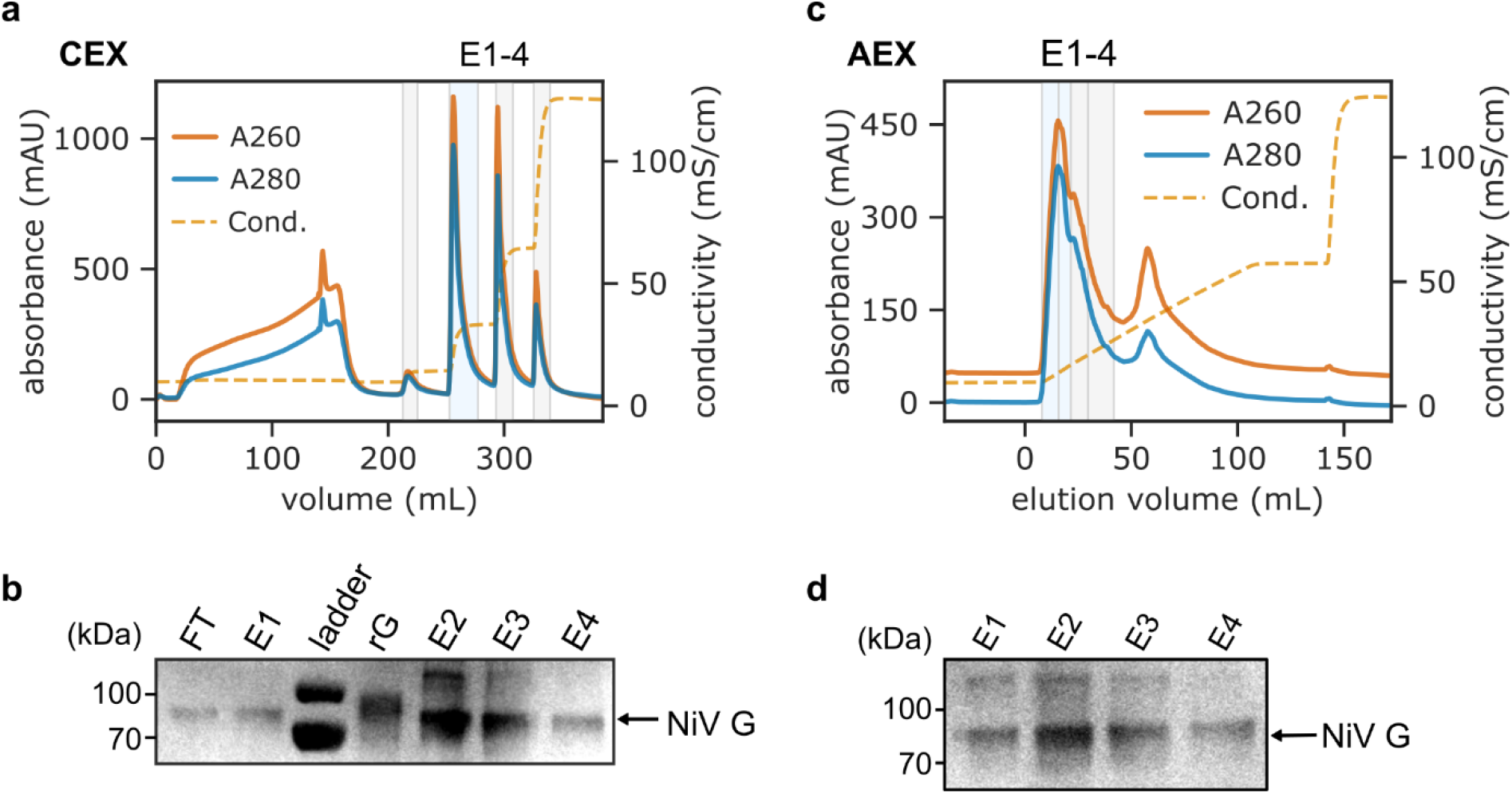
NiVLP purification with CEX and AEX chromatography. **(a)** Collected fractions from SXC were loaded on 8 mL CIM SO3 CEX column and purified fractionated using step gradient with increasing NaCl concentration. Shaded area denotes collected elution fractions E1-E4. E2, which was selected for further purification, is light blue. **(b)** NiVLP detection after CEX with western blot analysis against NiV G. The majority of NiV G is detected in E2 fraction, which was further purified on AEX. **(c)** The E2 fraction from CEX step was diluted and loaded on 4 mL CIM QA AEX column and separated in ascending salt gradient. **(d)** NiV G detection with western blot after AEX. The majority of NiVLPs are in E2 fraction. Fractions E1 and E2 were pooled and sterile filtered for immunogenicity study. FT: flow-through; E1-4: elution fractions; rG: recombinant NiV G.

The purified NiVLPs were sterile filtered and analyzed with NTA. The produced vaccine particles display 120 nm mean diameter, with 83 nm modal value as determined by NTA (Fig. 3a). This is in agreement with pleomorphic native NiV appearance (Hyatt *et al*., 2001; Ang, Lim and Wang, 2018). NTA measurements were also confirmed by TEM, which revealed characteristic enveloped particles with diameters around 100 nm and evidence of a protein corona, likely corresponding to NiV G and F glycoproteins (Fig. 3b). Thus, NiVLPs expressed in insect cells and purified by this multi-step monolith process closely mimic the morphology of the native pathogen. We used the sterile filtered NiVLPs to determine their immunogenicity in a Syrian hamster model.

**Fig. 3.**
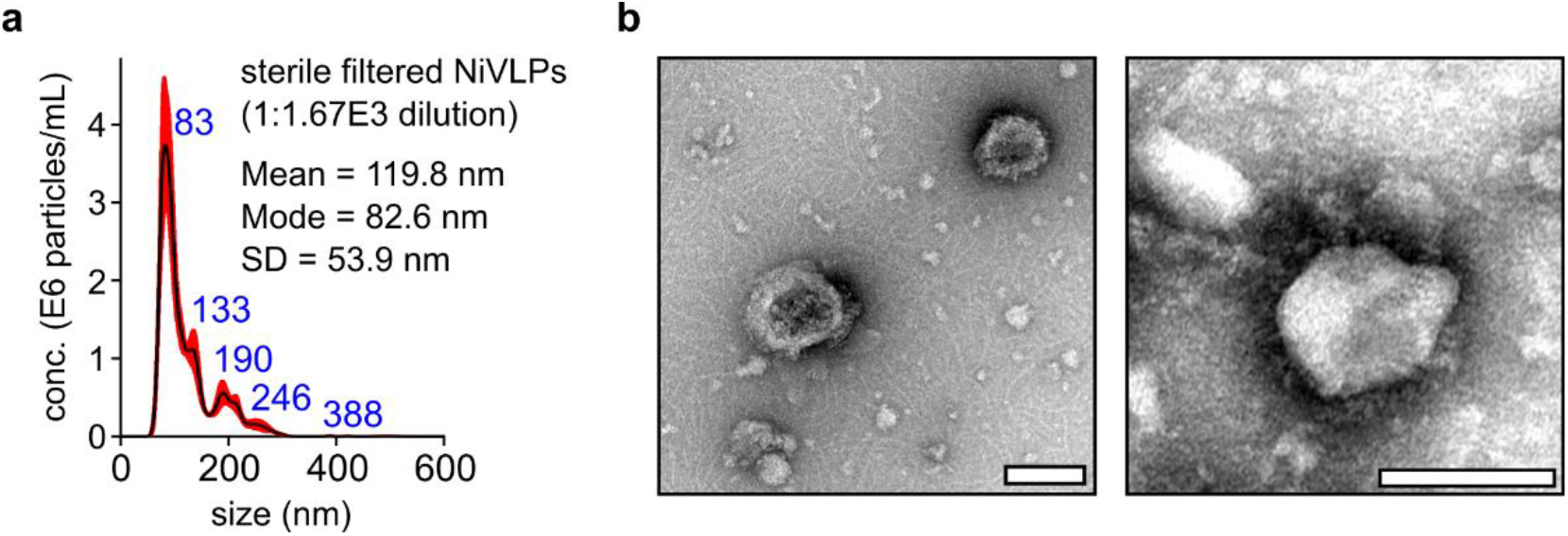
Purified NiVLP particles mimic native viral structure. **(a)** NTA size distribution analysis of sterile filtered NiVLPs. Red shaded area indicates standard error of the mean from 5 runs. SD: standard deviation. **(b)** TEM images of purified NiVLPs. Scale bars are 100 nm.

We immunized Syrian golden hamsters (n = 6) with 25 µg NiVLP intramuscularly (i.m.) with and without AddaVax (ADX) oil-in-water squalene adjuvant (Calabro *et al*., 2013) (Fig. 4a). Hamsters were boosted 21 days post-prime and we collected serum samples on days 0, 14, 28 and 42, when the study concluded. We quantified IgG titers against recombinant NiV G ectodomain (G_ecd_) with ELISA and present the data as area under the curve (AUC) (Fig. 4b). NiVLPs, both with and without adjuvant, induced a robust humoral response after a single dose. Interestingly, the ADX adjuvant did not significantly enhance IgG titers, even after the booster dose or at the study’s conclusion. We also determined the NiV-B neutralization potential of the final plasma samples in an in vitro neutralization assay (Fig. 5). Surprisingly, unadjuvanted NiVLPs generated slightly higher, though not statistically significant, neutralizing titers compared to the NiVLP + ADX formulation. However, serum from NiVLP-immunized animals (with or without adjuvant) exhibited low neutralizing activity against NiV-B, with only a few samples exceeding the limit of quantification (LOQ). Interestingly, NiVLPs expressed in mammalian cells induced higher levels of neutralizing antibodies after two doses and conferred complete protection against NiV challenge in hamsters after 1- and 3-dose regimen (Walpita *et al*., 2017). Altogether, while NiVLPs produced in insect cells are clearly immunogenic in terms of total IgG response, they generate low neutralizing antibody titers in the hamster model against NiV-B strain.

**Fig. 4.**
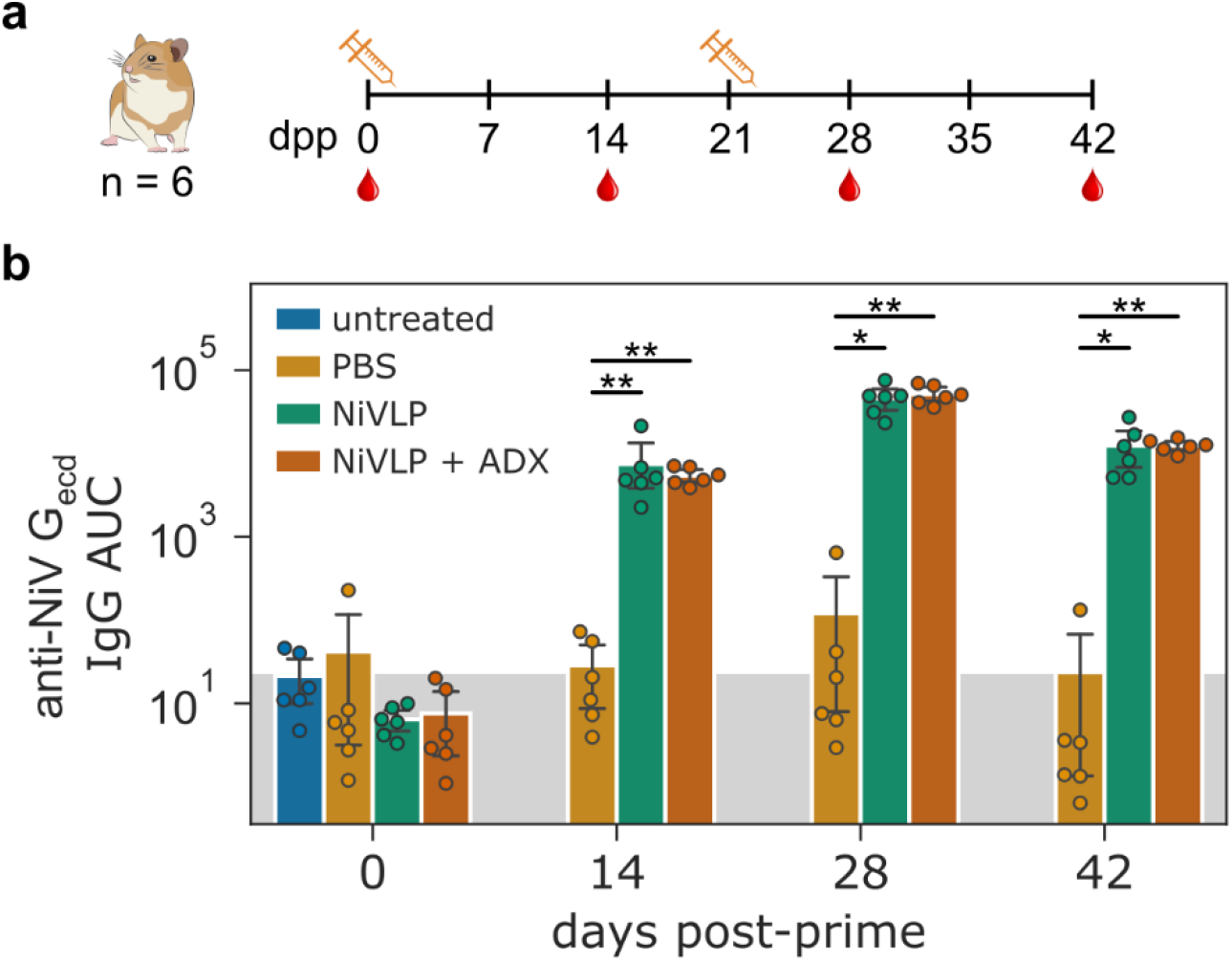
NiVLPs are immunogenic in hamsters after prime immunization. **(a)** NiVLP immunogenicity study plan. Syrian golden hamsters (n = 6) were immunized with 25 µg NiVLP with and without AddaVax adjuvant. Hamsters received booster dose 21 days post-prime. Sera samples were collected on days 0, 14, 28 and 42. dpp: days post-prime. **(b)** NiVLP immunogenicity was determined with anti-NiV G_ecd_ IgG ELISA. AUC: area under the curve; PBS: phosphate-buffered saline; ADX: AddaVax adjuvant. * p < 0.05, ** p < 0.01. For statistical comparison, log-transformed AUC titers were evaluated using a Kruskal-Wallis test corrected with Dunn’s multiple comparisons test.

**Fig. 5.**
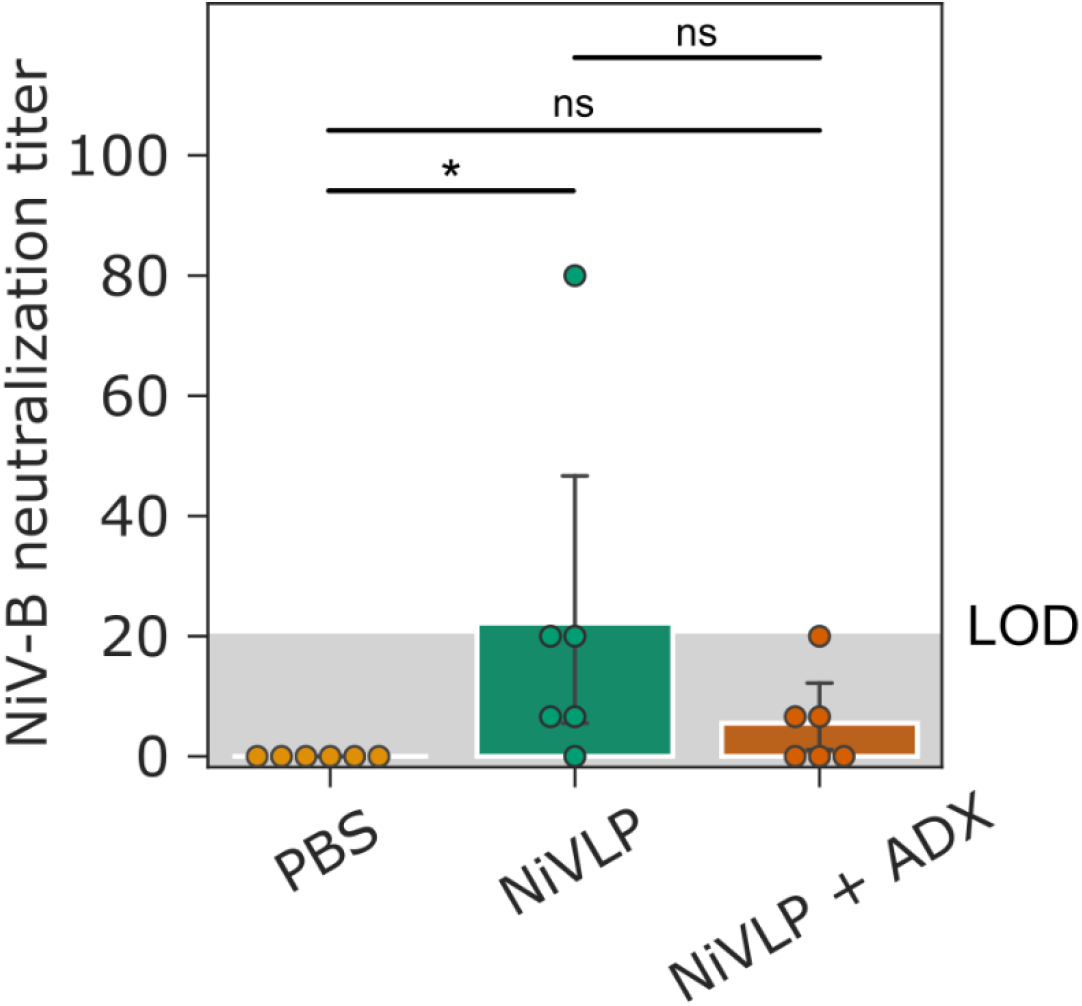
Limited NiVLPs stimulation of neutralizing immunity against NiV-B. Hamster serum samples 42 days post-prime immunization were used to determine neutralization titers against NiV-B in vitro. 50 TCID_50_ NiV-B was used for neutralization. PBS: phosphate-buffered saline; ADX: AddaVax adjuvant; LOD: limit of detection. * p < 0.05; ns: not significant. Statistical analysis was performed using Wilcoxon signed-rank test.

## Discussion

NiV poses a significant public health threat, yet no approved vaccine exists. VLPs offer a safe and effective vaccine strategy. While mammalian cell-derived NiVLPs are immunogenic (Walpita *et al*., 2011, Walpita *et al*., 2017; Rajan *et al*., 2024), their scalable production can be challenging. This study reports, to our knowledge, the first development of NiVLPs expressing F, G, and M proteins using a baculovirus system in Sf9 insect cells, coupled with an efficient, multi-step monolith chromatography purification.

Our scalable, chromatography-based purification strategy yielded highly pure NiVLPs of approximately 80-120 nm. The particle morphology was consistent with that of native NiV (Hyatt *et al*., 2001) and previously described NiVLPs (Walpita *et al*., 2011), suggesting correct assembly and surface glycoprotein display. The insect cell-derived NiVLPs induced robust immunogenicity in Syrian golden hamsters. A single 25 µg dose elicited strong systemic anti-NiV G_ecd_ IgG responses, even without adjuvant (Fig. 4b). This rapid induction of high antibody titers is a desirable characteristic for a potential outbreak vaccine. The oil-in-water adjuvant AddaVax (ADX) did not further enhance IgG levels, suggesting the particulate nature of the VLPs themselves is sufficiently immunostimulatory to drive a strong humoral response (Bachmann and Jennings, 2010).

Despite high total IgG titers, the neutralizing antibody response against the vaccine homolog Nipah virus Bangladesh strain (NiV-B) was low (Fig. 5). This finding contrasts with studies of mammalian cell-derived NiVLPs, which induced higher neutralization titers and protection in hamsters (Walpita *et al*., 2017). This discrepancy is likely attributable to differences in N-glycosylation patterns between insect (typically high-mannose) and mammalian cells (complex N-glycans) (Geisler and Jarvis, 2009). Glycosylation profoundly impacts viral glycoprotein folding and the presentation of antigens, including neutralizing epitopes. Altered G and F protein conformation on insect cell-derived NiVLPs could lead to limited neutralizing activity.

Interestingly, a structurally similar NiV replicon particle vaccine also failed to generate significant neutralizing titers in hamsters, yet was fully protective against NiV challenge (Welch *et al*., 2023). This challenges the key role of neutralizing antibody generation in Paramyxovirus vaccine efficacy determination (He *et al*., 2016; Pickering *et al*., 2016; Kingstad-Bakke *et al*., 2022). Therefore, a more detailed study of NiVLP cellular immune responses and Fc-dependent effector functions in relation to protection against Nipah infection is needed both in humans and in animal models (Bowman, Kaplonek and McNamara, 2023; Zhang *et al*., 2023).

In conclusion, this study establishes a scalable production and purification platform for morphologically correct NiVLPs that elicit potent systemic humoral immunity after a single dose. Future efforts could focus on evaluation of induced cellular immunity and protection against challenge, head-to-head comparison of insect cell and mammalian-derived antigens, utilizing mammalian-like glycoengineered insect cell lines for NiVLP production (Aumiller *et al*., 2012) and exploring alternative adjuvants to enhance neutralizing immunogenicity. Furthermore, a direct comparison of the antibody epitope specificity elicited by insect-versus mammalian-derived immunogens would be highly informative for multivalent NiVLP vaccine development.

## Materials and Methods

### Molecular Cloning and Recombinant Baculovirus Preparation

NiV-B G sequence was based on GenBank AEZ01397 sequence from 2010 Faridpur outbreak (Lo *et al*., 2012). NiV-M M sequence was based on GenBank AAF73379 sequence (Chua *et al*., 2000). NiV-M F sequence was based on GenBank AAK29087 with NiVop08 prefusion stabilization mutations (Loomis *et al*., 2020). The codon-optimized synthetic DNA constructs for insect cell expression were cloned into pLIB shuttle vector (Addgene 80610) in TOP10 *E. coli* cells (Invitrogen). The NiV M, F and G gene cassettes from pLIB plasmids were then combined into pBIG1a (Addgene 80611) vector using Gibson isothermal assembly according to the biGBac method (Weissmann *et al*., 2016). The expression cassettes were transferred into bacmid backbones in DH10Bac *E. coli* cells (Invitrogen). We isolated bacmid DNA with PureLink HiPure Plasmid Miniprep Kit (Invitrogen) and transfected Sf9 insect cells (Oxford Expression Technologies) using ExpiFectamine Sf reagent (Gibco) in a 6-well culture plate. Recombinant baculovirus (rBV) supernatants were harvested after 5–7 day incubation at 27 °C. We generated V1 baculovirus stocks by infecting Sf9 cells with 1:10 rBV supernatant dilution at 1·10^6^/mL. The viral stocks were harvested after incubation at 27 °C, 127 rpm for 3-4 days and stored at 4 °C. If necessary, V2 stocks were generated by infecting 2·10^6^/mL Sf9 cells with 1:100 V1 dilution. The rBV titers were determined with baculoQUANT qPCR quantification kit (Oxford Expression Technologies).

### Insect Cell Culture Expression and Harvest

Sf9 cells (Oxford Expression Technologies) were grown in a protein-free ESF 921 insect cell culture medium (Expression Systems) at 27 °C and 127 rpm. NiVLPs were expressed in 3 L polycarbonate vented shaker flasks (Corning) by infecting 1.2 L 1.5·10^6^/mL Sf9 cells at multiplicity of infection (MOI) 3. After 4 days of production the cell viability fell below 70 %, at which point we harvested culture supernatants by centrifugation at 4000 rcf for 5 min and 4 °C. The cell culture supernatant was stored at 4 °C prior NiVLP chromatography purification.

### NiVLP Chromatography Purification

NiVLP particles were purified from unfiltered insect cell culture supernatants. A 8 mL CIMmultus® OH column (6 µm; Sartorius BIA Separations) was equilibrated in 50 % buffer OH A (20 mM MES-NaOH pH 6.4, 500 mM NaCl, 7 % sucrose) and buffer OH B (20 mM MES-NaOH pH 6.4, 120 mM NaCl, 12 % w/v PEG-6000, 7 % sucrose), resulting in 6 % PEG-6000 concentration. For particle capture with SXC, we loaded the harvested samples on the CIMmultus® OH column with 1:1 in-line dilution with buffer OH B at 40 mL/min. We performed elution at 16 mL/min in a descending linear gradient from 50 % to 0 %B over 50 CV. The collected elution fractions were diluted 1:2 in 20 mM MES-NaOH pH 6.4, 7 % sucrose for nuclease digestion with DENARASE (c-LEcta) at 50 U/mL in presence of 2 mM MgCl_2_ at room temperature for 2 h to remove contaminating host cell nucleic acids. Digested samples were loaded at 40 mL/min on 8 mL CIMmultus® SO3 CEX column (6 µm; Sartorius BIA Separations), which was equilibrated with buffer SO3 A (20 mM MES-NaOH pH 6.4, 100 mM NaCl, 7 % sucrose). We performed elution in steps of 3-16-40-100 %B at 16 mL/min to buffer SO3 B (20 mM MES-NaOH pH 6.4, 2 M NaCl, 7 % sucrose). Selected fractions were diluted 1:2 with 20 mM HEPES-NaOH pH 7.4, 7 % sucrose and loaded at 20 mL/min on 4 mL CIMmultus® QA IEX column (6 µm; Sartorius BIA Separations), which was equilibrated with buffer QA A (20 mM HEPES-NaOH pH 7.4, 100 mM NaCl, 7 % sucrose). NiVLPs were isolated in linear elution gradient to 35 % buffer QA B (20 mM HEPES-NaOH pH 7.4, 2 M NaCl, 7 % sucrose) over 25 CVs. Finally, we exchanged buffer to PBS + 15 % sucrose with SEC group separation with Sepharose 6 Fast Flow resin on XK16/20 column (Cytiva) at 2 mL/min. NiVLPs were sterile filtered through 0.22 µm filters (TPP) and stored at −75 °C. The protein content was determined with Bio-Rad Protein Assay (Bio-Rad).

### SDS-PAGE and Western Blot

Collected samples were denatured at 70 °C for 10 min. We performed SDS-PAGE under reducing conditions on 4-20 % mPAGE™ Bis-Tris precast protein gels (Merck Millipore) in 1x MOPS running buffer at 200 V. We transferred the samples to activated Immobilion-P PVDF membrane (Merck Millipore) in a Mini-PROTEAN Trans-Blot Module (Bio-Rad) at 350 mA. Immunodetection was performed using SNAP i.d. system (Merck Millipore). The PVDF membrane was blocked with 0.5 % skim milk in TBST buffer. We used polyclonal anti-Nipah virus/HeV G protein primary antibodies (AntibodySystem, PVV07901). Secondary goat anti-rabbit HRP conjugates (Merck, 12-348) were used for colorimetric detection with 1-Step Ultra TMB-Blotting Solution (Thermo Scientific). Recombinant NiV G (Proteogenix, PX-P6271-100) protein was used as a positive control.

### Nanoparticle Tracking Analysis (NTA)

Particle size distribution and concentration were determined by Nanoparticle Tracking Analysis (NTA) using a NanoSight N300 (Malvern Instruments, United Kingdom) equipped with a blue laser module (488 nm). NTA software version 3.2 was used for capture and analysis of the data. NiVLP samples were diluted to a working concentration range of 10^7^–10^9^ particles/mL (20–100 particles per video frame) in particle-free water or storage buffer and injected into the device chamber using a syringe pump. Each sample was recorded five times for 45 s at constant capture settings (manually adjusted camera level and focus) and analysis parameters (detection threshold 4 or 5). All measurements were performed at room temperature.

### Transmission Electron Microscopy (TEM)

Formvar-coated grids were placed on drops of NiVLP samples for 5 min and negative staining was performed using 2% phosphotungstic acid (PTA). The grids were examined using a transmission electron microscope JEOL JEM-1400 Plus (Tokyo, Japan) at 120 kV.

### Hamster Immunogenicity Study

The hamster immunogenicity study was performed at the National Biosafety Laboratory, National Center for Public Health and Pharmacy, Budapest. The objective was to evaluate the immunogenicity of insect cell NiVLPs in golden Syrian hamsters. All animal experiments were performed according to the guidelines of the European Communities Council Directive (86/609 EEC) and were approved by the Hungarian National Authority under the license numbers of National Animal Ethical Commitee (PE/EA/00370-6/2024; PE/EA/00958-6/2024). The Syrian hamsters (Janvier) were 6 weeks old and kept under BSL-4 conditions in individually ventilated cages (IsoRat ISO48NFEEU, Techniplast, Italy), the body weight were measured weekly. Groups of 6 animals were assigned to either non-treated control, PBS-treated control, 25 µG NiVLP-only or 25 µg NiVLP + AddaVax™ (Invivogen) experimental groups. The blood samples were taken from orbital vein under inhalational anesthesia with isoflurane (isoflurane: ISOFLUTEK 1000 mg/g, Laboratorios Karizoo S.A.; anesthesia station: MiniHUB-V3, TEM SEGA, France) on dpp 0 (dpp: days post-prime), dpp 14 and dpp 28. At the end of the experiment on dpp 42, blood samples were taken via cardiac puncture after euthanasia. The non-treated control group was euthanized at dpp 0. For immunization 50 µL of NiVLPs were mixed 1:1 with AddaVax™ adjuvant or PBS (Gibco DPBS) and incubated for 15 minutes at room temperature and then used immediately. For PBS treated control group 100 µL PBS (Gibco DPBS) were used. All immunized hamsters received an intramuscular (i.m.) injection of 100 µL on dpp 0 and 21. The study was finished on day 42, all hamsters were euthanized and a final blood sample was collected via cardiac puncture. Serum was isolated from the blood and stored at −80° C.

### ELISA

IgG titers against NiV G were determined using an indirect ELISA protocol. Nunc-Immuno™ MicroWell™ 96 well plates (Sigma Aldrich) were coated overnight at 4 °C in 1xPBS with 0.5 µg/mL of recombinant NiV G_ecd_ (residues 172-602), which was produced in Sf9 insect cells and purified with Strep-tag affinity chromatography. Next, the plate was washed with wash buffer (0.01 M PBS pH 7.2, Tween-20 0.1% v/v) and blocked with 5% skim milk suspension for 1 h at room temperature. After washing, hamster serum samples were added to the plates at 1:50 initial dilution. Next, 5-fold serial dilutions were performed and plates were incubated at room temperature for 2 h. After washing, HRP-conjugated secondary goat anti-Syrian Hamster IgG H&L antibodies (Abcam, ab6892) at 1:10,000 dilution were added and incubated for 1 h at room temperature. The microtiter plates were incubated with TMB substrate (Sigma Aldrich, T4444) for 10 min. After, 2 M HCl stop solution was added and absorbance at 450 nm and 650 nm was determined. For area under the curve (AUC) titer calculation, we determined the baseline from mean blank values plus 3 standard deviations (SD). The negative values were set to 0. The AUC values were determined by integrating the serial dilution data with trapezoidal rule.

### In vitro Neutralization Assay

A seroneutralization assay was performed on hamster serum samples under BSL-4 conditions at the National Biosafety Laboratory, Budapest. Briefly, sera were inactivated at 56 °C for 30 min and were twofold serially diluted starting from 1:20 dilution using serum-free Dulbecco’s Modified Eagle’s Medium (DMEM, VWR) in triplicate in 96-well microtiter plates. Virus-only controls (diluted in DMEM) and cell-only controls (DMEM only) were run in parallel. An equal volume of Nipah virus Bangladesh strain (NiV-B), equivalent to 50 TCID_50_, was added to each well containing diluted sera and incubated for 1 hour at 37 °C. Subsequently, the mixtures were transferred onto 96-well cell culture plates (TPP, Switzerland) containing Vero E6 cells (Nuvonis) at approximately 2 · 10^5^ cells/mL in serum-free DMEM.

After 5 days of incubation at 37 °C and 5 % CO_2_, the plates were inactivated with 10 % formaldehyde solution and stained with crystal violet. The neutralizing antibody titer was defined as the reciprocal of the highest serum dilution that completely inhibited cytopathic effect. The limit of detection for this assay was 1:20 dilution. For calculation purposes, sera that demonstrated partial CPE inhibition but did not achieve complete neutralization were assigned a nominal titer of 6.6 for geometric mean determination.

## Statistical Analysis

The non-parametric Kruskal-Wallis statistical test with Dunn’s multiple comparisons test (Holm mehod) was performed with Python 3.7.11 using pandas, SciPy and scikit-posthocs packages. Asterisks denote correlations that had statistically significant p-value (* p < 0.05; ** p < 0.01).

## Availability of data and materials

All data generated or analyzed during this study are available from the corresponding author on reasonable request.

## Funding

This work benefited from access to the BSL4 laboratory at National Biosafety Laboratory, National Center for Public Health and Pharmacy, Budapest. The animal study and BSL-4 work are supported by the European Union’s Horizon Europe research and innovation program ISIDORE under grant agreement number 101046133 and received funding by the Slovenian Research and Innovation Agency (program No. P3-0083).

## Declaration of Competing Interest

UB and AJ are employees of Sferogen d.o.o., which is a vaccine development company. UB and MP are co-founders of Sferogen d.o.o. Other authors declare no competing interests.

## Authors’ contributions

UB conceptualized and wrote the manuscript, cloned the genetic constructs, performed chromatography purifications, ELISA and statistical analysis. AJ designed the hamster immunogenicity study and wrote the manuscript. TK cloned the genetic constructs, performed chromatography purifications and ELISA. MLK performed qPCR and western blot analytics. EB maintained insect cell culture, performed qPCR and western blot analytics. MK performed TEM. BP, KZ designed and performed the hamster immunogenicity study and seroneutralization assay. DD performed the hamster immunogenicity study and seroneutralization assay. MP conceptualized the study. All authors read and approved the final manuscript.

## Acknowledgements

We thank M. Leskovec and M. Šnajder of Sartorius BIA Separations for NTA analysis. The pLIB and pBIG1a plasmids were gifts from Jan-Michael Peters.

## References

Amaya, M. and Broder, C.C. (2020) ‘Vaccines to Emerging Viruses: Nipah and Hendra’, Annual Review of Virology, 7(1), pp. 447–473. Available at: 10.1146/annurev-virology-021920-113833.

Ang, B.S.P., Lim, T.C.C. and Wang, L. (2018) ‘Nipah Virus Infection’, Journal of Clinical Microbiology, 56(6), p. 10.1128/jcm.01875-17. Available at: 10.1128/jcm.01875-17.

Aumiller, J.J. et al. (2012) ‘A new glycoengineered insect cell line with an inducibly mammalianized protein N-glycosylation pathway’, Glycobiology, 22(3), pp. 417–428. Available at: 10.1093/glycob/cwr160.

Avanzato, V.A. et al. (2019) ‘A structural basis for antibody-mediated neutralization of Nipah virus reveals a site of vulnerability at the fusion glycoprotein apex’, Proceedings of the National Academy of Sciences, 116(50), pp. 25057–25067. Available at: 10.1073/pnas.1912503116.

Bachmann, M.F. and Jennings, G.T. (2010) ‘Vaccine delivery: a matter of size, geometry, kinetics and molecular patterns’, Nature Reviews Immunology, 10(11), pp. 787–796. Available at: 10.1038/nri2868.

Bezeljak, U. et al. (2025) ‘Development of multivalent SARS-CoV-2 virus-like particle vaccine candidates’, Vaccine, 61, p. 127394. Available at: 10.1016/j.vaccine.2025.127394.

Bowman, K.A., Kaplonek, P. and McNamara, R.P. (2023) ‘Understanding Fc function for rational vaccine design against pathogens’, mBio, 15(1), pp. e03036–23. Available at: 10.1128/mbio.03036-23.

Calabro, S. et al. (2013) ‘The adjuvant effect of MF59 is due to the oil-in-water emulsion formulation, none of the individual components induce a comparable adjuvant effect’, Vaccine, 31(33), pp. 3363– 3369. Available at: 10.1016/j.vaccine.2013.05.007.

Chua, K.B. et al. (2000) ‘Nipah virus: a recently emergent deadly paramyxovirus’, Science (New York, N.Y.), 288(5470), pp. 1432–1435. Available at: 10.1126/science.288.5470.1432.

Dang, H.V. et al. (2021) ‘Broadly neutralizing antibody cocktails targeting Nipah virus and Hendra virus fusion glycoproteins’, Nature Structural & Molecular Biology, 28(5), pp. 426–434. Available at: 10.1038/s41594-021-00584-8.

Donaldson, B. et al. (2018) ‘Virus-like particle vaccines: immunology and formulation for clinical translation.’, Expert review of vaccines, 17(9), pp. 833–849. Available at: 10.1080/14760584.2018.1516552.

van Doremalen, N. et al. (2022) ‘ChAdOx1 NiV vaccination protects against lethal Nipah Bangladesh virus infection in African green monkeys’, npj Vaccines, 7(1), pp. 1–8. Available at: 10.1038/s41541-022-00592-9.

Eaton, B.T. et al. (2006) ‘Hendra and Nipah viruses: different and dangerous’, Nature Reviews Microbiology, 4(1), pp. 23–35. Available at: 10.1038/nrmicro1323.

Effio, C.L. and Hubbuch, J. (2015) ‘Next generation vaccines and vectors: Designing downstream processes for recombinant protein-based virus-like particles’, Biotechnology Journal, 10(5), pp. 715– 727. Available at: 10.1002/biot.201400392.

Foster, S.L. et al. (2022) ‘A recombinant VSV-vectored vaccine rapidly protects nonhuman primates against lethal Nipah virus disease’, Proceedings of the National Academy of Sciences, 119(12), p. e2200065119. Available at: 10.1073/pnas.2200065119.

Gagnon, P. (2009) ‘Chromatographic Purification of Virus Particles’, in Encyclopedia of Industrial Biotechnology. Hoboken, NJ, USA: John Wiley & Sons, Inc. Available at: 10.1002/9780470054581.eib583.

Gao, Z. et al. (2022) ‘Assessment of the immunogenicity and protection of a Nipah virus soluble G vaccine candidate in mice and pigs’, Frontiers in Microbiology, 13, p. 1031523. Available at: 10.3389/fmicb.2022.1031523.

Geisbert, T.W. et al. (2021) ‘A single dose investigational subunit vaccine for human use against Nipah virus and Hendra virus’, NPJ Vaccines, 6, p. 23. Available at: 10.1038/s41541-021-00284-w.

Geisler, C. and Jarvis, D. (2009) ‘Insect Cell Glycosylation Patterns in the Context of Biopharmaceuticals’, Post-translational Modification of Protein Biopharmaceuticals, pp. 165–191. Available at: 10.1002/9783527626601.ch7.

He, W. et al. (2016) ‘Epitope specificity plays a critical role in regulating antibody-dependent cell-mediated cytotoxicity against influenza A virus’, Proceedings of the National Academy of Sciences, 113(42), pp. 11931–11936. Available at: 10.1073/pnas.1609316113.

Hyatt, A.D. et al. (2001) ‘Ultrastructure of Hendra virus and Nipah virus within cultured cells and host animals’, Microbes and Infection, 3(4), pp. 297–306. Available at: 10.1016/S1286-4579(01)01383-1.

Ithinji, D.G. et al. (2022) ‘Multivalent viral particles elicit safe and efficient immunoprotection against Nipah Hendra and Ebola viruses’, NPJ Vaccines, 7. Available at: 10.1038/s41541-022-00588-5.

Ker, D.-S. et al. (2021) ‘CryoEM structure of the Nipah virus nucleocapsid assembly’, PLOS Pathogens, 17(7), p. e1009740. Available at: 10.1371/journal.ppat.1009740.

Khan, S. et al. (2024) ‘Twenty-five years of Nipah outbreaks in Southeast Asia: A persistent threat to global health’, IJID Regions, 13, p. 100434. Available at: 10.1016/j.ijregi.2024.100434.

Kim, S. et al. (2025) ‘Progress and challenges in Nipah vaccine development and licensure for epidemic preparedness and response’, Expert Review of Vaccines, 24(1), pp. 183–193. Available at: 10.1080/14760584.2025.2476523.

Kingstad-Bakke, B. et al. (2022) ‘Vaccine-induced systemic and mucosal T cell immunity to SARS-CoV-2 viral variants’, Proceedings of the National Academy of Sciences, 119(20), p. e2118312119. Available at: 10.1073/pnas.2118312119.

Larsen, B.B. et al. (2025) ‘Functional and antigenic landscape of the Nipah virus receptor-binding protein’, Cell [Preprint]. Available at: 10.1016/j.cell.2025.02.030.

Lee, J. et al. (2012) ‘Principles and applications of steric exclusion chromatography’, Journal of Chromatography A, 1270, pp. 162–170. Available at: 10.1016/j.chroma.2012.10.062.

Lo, M.K. et al. (2012) ‘Characterization of Nipah virus from outbreaks in Bangladesh, 2008-2010’, Emerging Infectious Diseases, 18(2), pp. 248–255. Available at: 10.3201/eid1802.111492.

Lo, M.K. et al. (2020) ‘Evaluation of a Single-Dose Nucleoside-Modified Messenger RNA Vaccine Encoding Hendra Virus-Soluble Glycoprotein Against Lethal Nipah virus Challenge in Syrian Hamsters’, The Journal of Infectious Diseases, 221(Supplement_4), pp. S493–S498. Available at: 10.1093/infdis/jiz553.

Loomis, R.J. et al. (2020) ‘Structure-Based Design of Nipah Virus Vaccines: A Generalizable Approach to Paramyxovirus Immunogen Development’, Frontiers in Immunology, 11. Available at: https://www.frontiersin.org/articles/10.3389/fimmu.2020.00842 (Accessed: 11 November 2022).

Loomis, R.J. et al. (2021) ‘Chimeric Fusion (F) and Attachment (G) Glycoprotein Antigen Delivery by mRNA as a Candidate Nipah Vaccine’, Frontiers in Immunology, 12, p. 772864. Available at: 10.3389/fimmu.2021.772864.

Lu, M. et al. (2023) ‘Both chimpanzee adenovirus-vectored and DNA vaccines induced long-term immunity against Nipah virus infection’, npj Vaccines, 8(1), pp. 1–12. Available at: 10.1038/s41541-023-00762-3.

Margine, I. et al. (2012) ‘Residual Baculovirus in Insect Cell-Derived Influenza Virus-Like Particle Preparations Enhances Immunogenicity’, PLOS ONE, 7(12), p. e51559. Available at: 10.1371/journal.pone.0051559.

Minh, A. and Kamen, A.A. (2021) ‘Critical Assessment of Purification and Analytical Technologies for Enveloped Viral Vector and Vaccine Processing and Their Current Limitations in Resolving Co-Expressed Extracellular Vesicles’, Vaccines, 9(8), p. 823. Available at: 10.3390/vaccines9080823.

Moore, K.A. et al. (2024) ‘Measures to prevent and treat Nipah virus disease: research priorities for 2024–29’, The Lancet Infectious Diseases, 24(11), pp. e707–e717. Available at: 10.1016/S1473-3099(24)00262-7.

Morenweiser, R. (2005) ‘Downstream processing of viral vectors and vaccines’, Gene Therapy, 12(1), pp. S103–S110. Available at: 10.1038/sj.gt.3302624.

Mortola, E. and Roy, P. (2004) ‘Efficient assembly and release of SARS coronavirus-like particles by a heterologous expression system’, FEBS Letters, 576(1–2), pp. 174–178. Available at: 10.1016/j.febslet.2004.09.009.

Mungall, B.A. et al. (2006) ‘Feline Model of Acute Nipah Virus Infection and Protection with a Soluble Glycoprotein-Based Subunit Vaccine’, Journal of Virology, 80(24), pp. 12293–12302. Available at: 10.1128/JVI.01619-06.

Nestola, P. et al. (2015) ‘Improved virus purification processes for vaccines and gene therapy.’, Biotechnology and bioengineering, 112(5), pp. 843–57. Available at: 10.1002/bit.25545.

Noad, R. and Roy, P. (2003) ‘Virus-like particles as immunogens’, Trends in Microbiology, 11(9), pp. 438–444. Available at: 10.1016/S0966-842X(03)00208-7.

Nooraei, S. et al. (2021) ‘Virus-like particles: preparation, immunogenicity and their roles as nanovaccines and drug nanocarriers’, Journal of Nanobiotechnology, 19(1), p. 59. Available at: 10.1186/s12951-021-00806-7.

Pallister, J.A. et al. (2013) ‘Vaccination of ferrets with a recombinant G glycoprotein subunit vaccine provides protection against Nipah virus disease for over 12 months’, Virology Journal, 10, p. 237. Available at: 10.1186/1743-422X-10-237.

Pedrera, M. et al. (2024) ‘Evaluation of the immunogenicity of an mRNA vectored Nipah virus vaccine candidate in pigs’, Frontiers in Immunology, 15. Available at: 10.3389/fimmu.2024.1384417.

Pickering, B.S. et al. (2016) ‘Protection against henipaviruses in swine requires both, cell-mediated and humoral immune response’, Vaccine, 34(40), pp. 4777–4786. Available at: 10.1016/j.vaccine.2016.08.028.

Rajan, A. et al. (2024) ‘Highly sensitive and quantitative HiBiT-tagged Nipah virus-like particles: A platform for rapid antibody neutralization studies’, Heliyon, 10(11), p. e31905. Available at: 10.1016/j.heliyon.2024.e31905.

Smith, G.E. et al. (2013) ‘Development of influenza H7N9 virus like particle (VLP) vaccine: Homologous A/Anhui/1/2013 (H7N9) protection and heterologous A/chicken/Jalisco/CPA1/2012 (H7N3) cross-protection in vaccinated mice challenged with H7N9 virus’, Vaccine, 31(40), pp. 4305–4313. Available at: 10.1016/j.vaccine.2013.07.043.

Walpita, P. et al. (2011) ‘Vaccine Potential of Nipah Virus-Like Particles’, PLOS ONE, 6(4), p. e18437. Available at: 10.1371/journal.pone.0018437.

Walpita, P. et al. (2017) ‘A VLP-based vaccine provides complete protection against Nipah virus challenge following multiple-dose or single-dose vaccination schedules in a hamster model’, npj Vaccines, 2(1), pp. 1–9. Available at: 10.1038/s41541-017-0023-7.

Watkinson, R.E. and Lee, B. (2016) ‘Nipah virus matrix protein: expert hacker of cellular machines’, FEBS Letters, 590(15), pp. 2494–2511. Available at: 10.1002/1873-3468.12272.

Weissmann, F. et al. (2016) ‘BiGBac enables rapid gene assembly for the expression of large multisubunit protein complexes’, Proceedings of the National Academy of Sciences of the United States of America, 113(19), pp. E2564–E2569. Available at: 10.1073/pnas.1604935113.

Welch, S.R. et al. (2023) ‘Single-dose mucosal replicon-particle vaccine protects against lethal Nipah virus infection up to 3 days after vaccination’, Science Advances, 9(31), p. eadh4057. Available at: 10.1126/sciadv.adh4057.

de Wit, E. et al. (2022) ‘Distinct VSV-based Nipah virus vaccines expressing either glycoprotein G or fusion protein F provide homologous and heterologous protection in a nonhuman primate model’, eBioMedicine, 87, p. 104405. Available at: 10.1016/j.ebiom.2022.104405.

Wit, E. de et al. (2023) ‘Distinct VSV-based Nipah virus vaccines expressing either glycoprotein G or fusion protein F provide homologous and heterologous protection in a nonhuman primate model’, eBioMedicine, 87. Available at: 10.1016/j.ebiom.2022.104405.

Wong, J.J. et al. (2021) ‘EphrinB2 clustering by Nipah virus G is required to activate and trap F intermediates at supported lipid bilayer–cell interfaces’, Science Advances, 7(5), p. eabe1235. Available at: 10.1126/sciadv.abe1235.

Xu, K. et al. (2008) ‘Host cell recognition by the henipaviruses: Crystal structures of the Nipah G attachment glycoprotein and its complex with ephrin-B3’, Proceedings of the National Academy of Sciences of the United States of America, 105(29), pp. 9953–9958. Available at: 10.1073/pnas.0804797105.

Yoneda, M. et al. (2013) ‘Recombinant Measles Virus Vaccine Expressing the Nipah Virus Glycoprotein Protects against Lethal Nipah Virus Challenge’, PLOS ONE, 8(3), p. e58414. Available at: 10.1371/journal.pone.0058414.

Zhang, A. et al. (2023) ‘Beyond neutralization: Fc-dependent antibody effector functions in SARS-CoV-2 infection’, Nature Reviews Immunology, 23(6), pp. 381–396. Available at: 10.1038/s41577-022-00813-1.

